# Process for Standardizing and Assessing the Parameters Governing MS2 Virus-Like Particle Reassembly around Nucleic Acid Cargo

**DOI:** 10.64898/2025.12.02.691839

**Authors:** Daniel de Castro Assumpcao, Emma S. Vinokour, Madeline M. Mills, Shiqi Liang, Carolyn E. Mills, Aline Carvalho da Costa, Nolan W. Kennedy, Danielle Tullman-Ercek

**Affiliations:** Department of Chemical and Biological Engineering, Northwestern University, Evanston, Illinois, USA; Weinberg College of Arts and Sciences, Northwestern University, Evanston, Illinois, USA; Interdisciplinary Biological Sciences Program, Northwestern University, Evanston, Illinois, USA; Department of Bioengineering, University of California, Santa Barbara, California, USA; School of Chemical Engineering, University of Campinas, Campinas, Sao Paulo, Brazil

**Keywords:** Capsid assembly, design of experiments, self-assembly

## Abstract

MS2 virus-like particles (VLPs) are widely used as protein nanocages for cargo encapsulation, yet *in vitro* disassembly–reassembly protocols remain poorly standardized, and reassembly yields are reported inconsistently. As a result, the same experiments reported in literature produce widely divergent yields, limiting reproducibility and cross-study comparability. Here, we introduce a cargo-specific, quantitative framework for standardized MS2 VLP reassembly yield determination. We evaluate commonly used disassembly and post-disassembly processing methods and identify practical trade-offs between protein recovery, accessibility, and reproducibility. Reassembly yield is quantified using size exclusion chromatography calibrated against purified VLP standards, enabling robust, cargo-specific yield measurement. Using this framework, we apply a full factorial design of experiments to quantify the individual and combined effects of coat protein concentration, ionic strength, buffer pH, and molecular crowding on reassembly yield. The resulting statistical model explains more than 99% of the explainable variance and its linear fit to the experimental data indicates that optimal reassembly conditions extend beyond those tested to date. Protein concentration and ionic strength dominate reassembly yield, whereas pH and osmolyte concentration contribute more modestly within the tested ranges. Finally, we propose practical guidelines for standardized MS2 VLP disassembly, reassembly, and yield reporting, defining a transferable operating envelope for MS2 VLP reconstruction. While demonstrated here using a single nucleic acid cargo (tr-DNA), the framework is readily extensible to alternative cargos and coat protein variants.

## INTRODUCTION

The MS2 virus-like particle (VLP) is a protein-based nanoparticle derived from the capsid of the male-specific bacteriophage MS2 (*Emesvirus zinderi*) [1]. Composed solely of 180 identical copies of a 129-amino acid <u>c</u>oat <u>p</u>rotein (CP), the MS2 VLP assembles into a 27 nm icosahedral structure that lacks viral RNA and is non-infectious [2,3]. These protein nanoparticles can be readily produced in *Escherichia coli* using plasmid-based expression systems, where the expressed CPs spontaneously self-assemble around RNA molecules, including their encoding mRNA during translation [4]. Owing to their simplicity, symmetry, and genetic tractability, MS2 VLPs have become a widely used model in virology and a versatile platform in nanotechnology. They have been extensively investigated for applications in imaging, diagnostics, vaccination, and targeted drug delivery [5–7]. In therapeutic delivery, a common strategy involves disassembling the capsid to remove encapsulated bacterial RNA (“native” cargo), followed by reassembly in a buffered solution containing the cargo of interest resulting in a protein-based nanoparticle loaded with a desired molecule [8–17]. Despite the widespread use of this approach, experimental protocols remain highly heterogeneous, and researchers seldom report standardized metrics for evaluating reassembly efficiency. As a result, nominally identical MS2 disassembly–reassembly experiments have reported widely divergent yields, and a standardized method for MS2 disassembly/reassembly, as well as reproducible and quantitative reassembly yield estimation, have yet to be formalized.

Some alternative strategies avoid disassembly altogether, relying instead on *in vivo* expression systems to alter the cargo [18], *in vivo* cargo conjugation to the interior of the VLP [19], or cargo replacement without disassembling the VLP [20]. Nevertheless, the predominant *in vitro* protocol for cargo loading involves four main steps: (1) disassembling the MS2 VLP using acidic conditions to generate free coat protein, (2) removing the encapsulated native mRNA, (3) adjusting the pH of the coat protein solution, and (4) incubating the coat proteins with the target cargo under reassembly conditions (**Figure 1**). Despite the widespread use of this general framework, the specific conditions used in each step vary widely across studies. For example, incubation time and acid concentration during disassembly differ significantly (e.g. 30 min to 2 h), and reassembly conditions, such as buffer composition, pH, and the inclusion of crowding agents or salts, also lack standardization. Moreover, few publications clearly define, quantify, or report reassembly yield in a consistent manner, making it difficult to compare reassembly efficiencies across different protocols or draw conclusions from the existing literature (see **Supplemental Information 1** for an extensive comparison between experimental conditions across literature).

**Figure 1:**
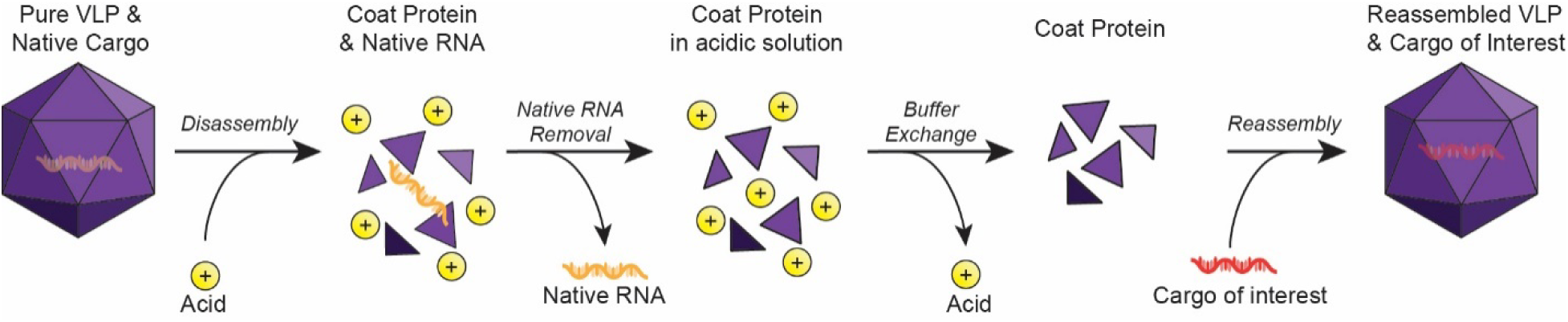
Overall process for encapsulating a cargo of interest in MS2 VLPs. Biologically produced and purified MS2 VLPs are first incubated under acidic conditions to induce capsid disassembly and precipitate nucleic acids natively packed during VLP expression. The precipitated nucleic acids are removed by centrifugation, and the resulting coat protein solution is buffer exchanged to eliminate excess acid and adjust the pH. Finally, the coat proteins are incubated with the cargo of interest in buffer, promoting spontaneous self-assembly and encapsulation of cargo with total or local negative charges on its structure.

In this study, we systematically investigate each step of the *in vitro* cargo-loading process for MS2 VLPs, with a focus on the protein-level parameters and solution conditions that influence disassembly and reassembly. First, we examine how variations in acid incubation time affect the removal of native RNA. We then discuss alternative methods for adjusting the pH of the coat protein solution post-disassembly. Next, we evaluate current approaches for calculating reassembly yield and propose a quantitative, reproducible metric designed to enable cross-study comparability. Finally, we implement a full factorial <u>d</u>esign <u>o</u>f <u>e</u>xperiments (DOE) framework to simultaneously assess the effects of buffer pH, ionic strength, protein concentration, and molecular crowding on reassembly efficiency. Through this comprehensive analysis, we define a method for MS2 VLP disassembly–reassembly, highlight the strengths and limitations of commonly used methods and propose standardized conditions for MS2 VLP disassembly and reassembly. While this study focuses on a single nucleic acid cargo (tr-DNA), the framework introduced here is designed to be extensible to alternative cargos and coat protein variants, providing a transferable foundation for systematic and reproducible MS2 VLP engineering.

## RESULTS AND DISCUSSION

### Incubation time with acetic acid impacts native cargo removal during disassembly

The first step in loading non-native cargo into MS2 VLPs involves disassembling the capsid and removing encapsulated native RNA using glacial acetic acid. Here, the acid serves two purposes: to disassemble the VLP and to precipitate its native nucleic acid cargo. Most published protocols employ a 2:1 volumetric ratio of glacial acetic acid to VLP solutions with VLPs typically at 10 mg/mL [8,10–17,21]. However, some studies report using a 3:1 ratio [9] or lower VLP concentrations (e.g., 4 mg/mL) [16]. Among the reported parameters, incubation time varies the most significantly across literature, from 30 min [8,10,12,14–17] to 2 h [9] (See **Supplemental Information 1** for a comprehensive comparison). To identify optimal conditions for VLP disassembly, we considered the three primary goals of this step: (1) complete dissociation of the capsid into coat protein subunits, (2) efficient precipitation and removal of the encapsulated native RNA, and (3) maximum possible coat protein concentration after disassembly. This last goal (maximum coat protein concentration) enables the choice of a wider range of initial coat protein concentrations in reassembly reactions.

To determine whether the lowest reported acetic acid-to-VLP ratio (2:1 v/v) and the highest reported VLP concentration (10 mg/mL) are sufficient for complete capsid disassembly, we analyzed samples before acid treatment and after acid treatment followed by centrifugation using <u>s</u>ize-<u>e</u>xclusion <u>c</u>hromatography (SEC) and transmission <u>e</u>lectron <u>m</u>icroscopy (TEM) (**Figure 2A–B**). The shift in SEC retention time from 8.1 min (intact capsid) to 14.0 min (disassembled protein) and the absence of intact particles in TEM micrographs indicate that these conditions fully disassemble the capsids. We confirmed protein integrity preservation after disassembly with sodium dodecyl sulfate–polyacrylamide gel electrophoresis (SDS–PAGE) (**Figure 2C**).

**Figure 2:**
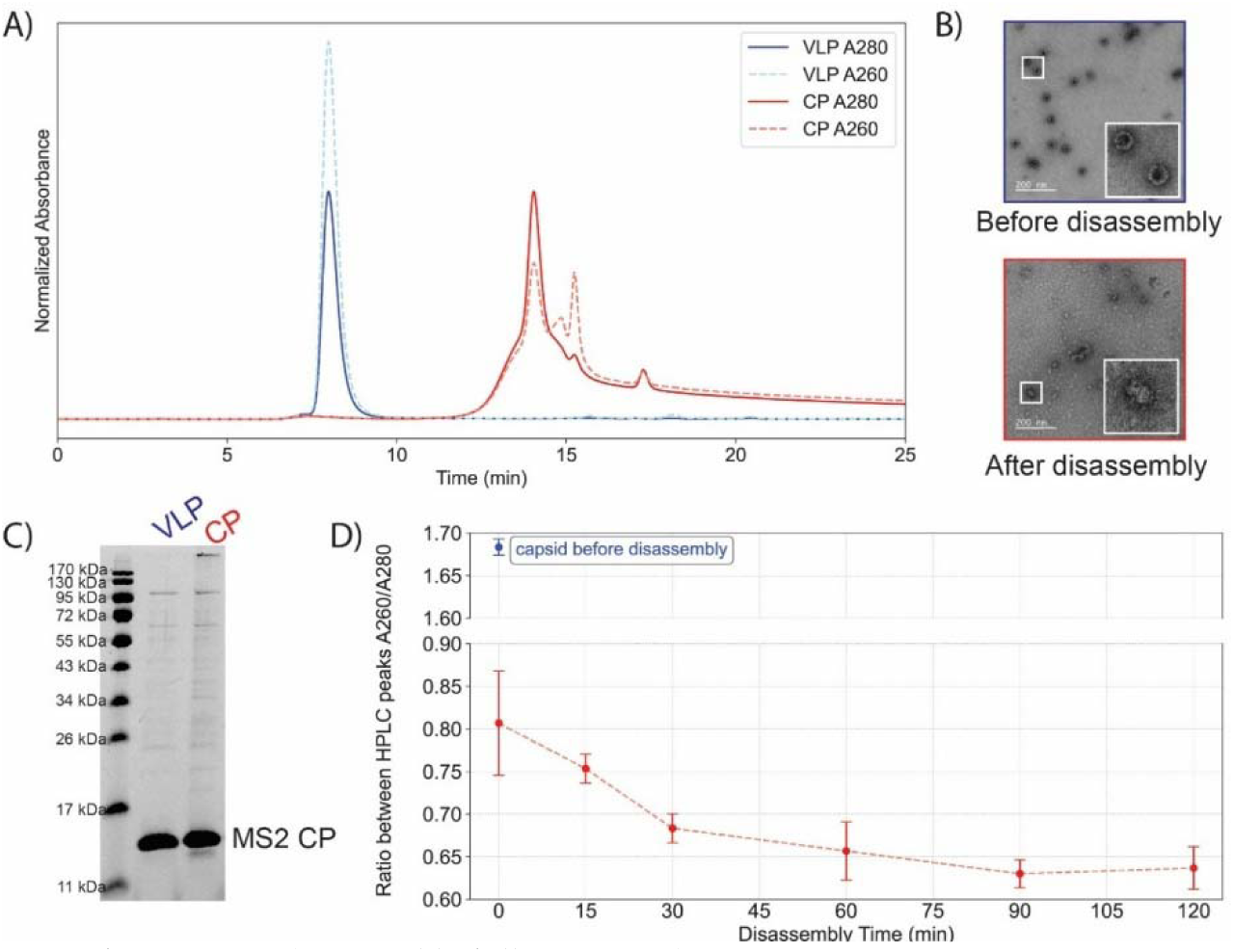
Characterization of MS2 VLP disassembly following acid treatment. A) Size exclusion chromatography (SEC) traces showing a shift in retention time before (labelled “VLP”) and after (labelled “CP”) incubation with acetic acid, indicating capsid disassembly. A concurrent decrease in A□□□ absorbance reflects the removal of native nucleic acid via precipitation and centrifugation. B) Transmission electron microscopy (TEM) images of VLPs before and after acid incubation, demonstrating the transition from intact capsids to irregular protein aggregates. C) SDS-PAGE of samples before and after disassembly in acetic acid indicating that protein does not degrade during disassembly D) A□□□/A□□□ absorbance ratio of the protein solution as a function of incubation time, showing that RNA removal improves with longer incubation and reaches a plateau after ∼90 min.

In addition to disassembling the capsid, the low pH generated by glacial acetic acid also promotes spontaneous precipitation of the native RNA packaged within MS2 VLPs. Efficient removal of this RNA is a critical step in the disassembly–reassembly process, as residual negatively charged nucleic acids can trigger unintended, premature self-assembly during pH neutralization, prior to the introduction of the desired cargo. A simple diagnostic metric to confirm the removal of nucleic acids from VLP is the <u>u</u>ltra<u>v</u>iolet-<u>vis</u>ible spectroscopy (UV-Vis) absorbance profiles, particularly ratio between absorbances at 260 nm and 280 nm. Pure RNA has an A_260_/A_280_ of approximately 2.1 and pure proteins have an A_260_/A_280_ of approximately 0.6. These values are heuristic with widespread use in the field, rather than a theoretical value and thus, may change according to the molecule composition. For MS2, we observe a drop in the A_260_/A_280_ ratio of samples pre-disassembly (A_260_/A_280_ ∼1.65) and post-disassembly followed by centrifugation (A_260_/A_280_ ∼0.65), suggesting removal of the native cargo (**Figure 2A**). As noted by Garmann *et al.* [11], the clarified coat protein solution should ideally have an A_260_/A_280_ ratio below 0.67 — preferably closer to 0.60 — to confirm effective removal of RNA.

Nucleic acid precipitation is driven by its protonation under acidic conditions, which decreases its solubility and enables nucleic acid separation from protein through standard solid–liquid separation techniques. While a single step centrifugation is used by all the consulted references, there is no consensus in the operational conditions (**Supplemental Information 1**). We tested only one condition, in which centrifugation for 15 min at 17,000 × *g* and 4 °C was sufficient to pellet nucleic acids and achieve an A_260_/A_280_ = 0.65. Centrifugation followed by filtration could also be employed if the filter material is compatible with glacial acetic acid. Chromatography should be avoided as passing insoluble material through a column can cause clogging and reduce column lifespan.

To assess the impact of VLP-acetic acid incubation time on RNA removal, we measured A_260_/A_280_ values following disassembly of 10 mg/mL VLPs in a 2:1 acetic acid solution across varying incubation durations. Our results show that a 90-min incubation consistently produces coat protein solutions with A_260_/A_280_ ratios below 0.65, suggesting efficient RNA removal and optimal conditions for downstream reassembly (**Figure 2D**). Since not all laboratories have access to identical equipment, acid incubation time and centrifugation operational conditions should be tailored to specific experimental setups to ensure that this ratio approaches the ideal value of 0.60. It is also important to note that very high protein concentrations (>30 mg/mL) may lead to coat protein loss due to aggregation and should be tested prior to establishing internal protocols.

Once the capsid is disassembled and the native nucleic acid is removed, the next step is to eliminate the excess acetic acid. Common methods reported in the literature include desalting columns [8,12,14,15,17], dialysis [9,13,16], and centrifugal filtration [10] (See **Supplemental Information 1** for operational details), each with distinct advantages and limitations. Centrifugal filtration is the fastest and most cost-effective approach but is also labor-intensive and can promote premature protein aggregation, especially if local protein concentrations become high near the filter membrane. Desalting columns offer an alternative approach and can be operated via either high-performance or gravity flow liquid chromatography. While high-performance liquid chromatography allows for higher precision, it requires capital equipment that may not be accessible in all settings. Gravity-based desalting is simple and accessible but can lead to protein loss due to residual acidity, as some collected fractions may retain low pH values due to the non-automated nature of the technique and thus should be excluded from reassembly steps. Dialysis offers the highest protein recovery at a reassembly-compatible pH, is straightforward to perform, and is inexpensive; however, it is the most time-consuming method. Overall, each of these approaches is acceptable for acetic acid removal, and the optimal choice depends on laboratory constraints such as available resources, equipment, and desired throughput. In this work, we adopted dialysis to maximize protein recovery and ensure precise pH control of the coat protein solution prior to reassembly.

Another critical variable in this buffer exchange step is the choice of buffer. An effective buffer should maintain the coat proteins in a disassembled and soluble state, prevent precipitation, and avoid introducing components that interfere with downstream reassembly. Most studies recommend acetic acid solutions ranging from 1 mM (pH ∼4.4) to 20 mM (pH ∼3.7) (See **Supplemental Information 1** for additional buffer compositions). For example, Ashley *et al.* [9] employed a 10 mM acetic acid buffer supplemented with 50 mM sodium chloride. While sodium chloride can enhance protein solubility, its effect on disassembled coat proteins remains unclear. Excess ionic strength may promote premature capsid reassembly in the absence of cargo, thereby reducing encapsulation efficiency. For this reason, we use a 1 mM acetic acid solution with no salts as a dialysis buffer. The impact of salt concentration on reassembly efficiency is addressed in detail below.

### Methods for determining reassembly yield and cargo impact on quantification

Before reassembly conditions can be optimized, reliable and standardized methods for quantifying reassembly yield must be established. Because reassembly rarely reaches 100% efficiency, the final mixture inevitably contains both assembled VLPs and unassembled coat protein (CP). Quantification of reassembly yield therefore requires two steps: (1) separation of assembled VLPs from free CP, and (2) determination of the protein concentrations in the VLP fraction and the total solution before the reassembly reaction. Among studies that report yield of reassembly, the most common approaches for this quantification are colorimetric protein assays such as bicinchoninic acid (BCA) assay; densitometric analysis of protein bands resolved by SDS–PAGE; and absorbance-based techniques such as UV–vis spectroscopy, coupled or not to liquid chromatography (*i.e.* SEC) instruments (see **Supplemental Information 1** for methods details).

The BCA assay is a colorimetric, protein-specific method that quantifies protein concentration based on the reduction of Cu² to Cu by peptide bonds and thus not affected by the presence of nucleic acids. BCA assay works well for quantifying total protein. However, it is unsuitable for determining true reassembly yield if VLP reassembly is done around another encapsulated protein because it cannot distinguish between the two. We also considered densitometric analysis of SDS–PAGE bands, a semi-quantitative approach that correlates band intensity with protein concentration. Like BCA, this method is also not confounded by nucleic acids and can therefore provide informative estimates of reassembly yield when protein loading in the gel is carefully controlled. However, its utility is limited if the cargo is a protein of similar molecular weight to the MS2 coat protein. SEC appears particularly advantageous for VLPs because it simultaneously separates assembled from unassembled particles while providing a measurement (A_280_ peak area) that correlates to capsid concentration, seemingly condensing two processes into a single procedure. However, as is true for all absorbance-based methods, the absorbance at 280 nm is influenced by cargo composition. We demonstrated this by reassembling MS2 around either luciferase RNA (∼1900 nt) or a short tr-DNA (19 nt). Following nuclease digestion of unencapsulated cargo and SEC purification, both samples were brought to equal protein concentration as determined by BCA assay. When analyzed by SEC (**Figure 3A**), the luciferase RNA sample (A_260_/A_280_ = 1.45) produced a stronger A_280_ signal than the tr-DNA sample (A_260_/A_280_ = 1.09), despite identical protein concentrations, illustrating how cargo composition can confound absorbance-based yield quantification. In the case of this procedure, estimating the yield by comparing the absorbance of unassembled capsid with reassembled capsid would overestimate yield. Therefore, to establish a reliable correlation between A_280_ peak area and protein concentration, we recommend using the reassembled product itself as a calibrant, described in detail below.

**Figure 3:**
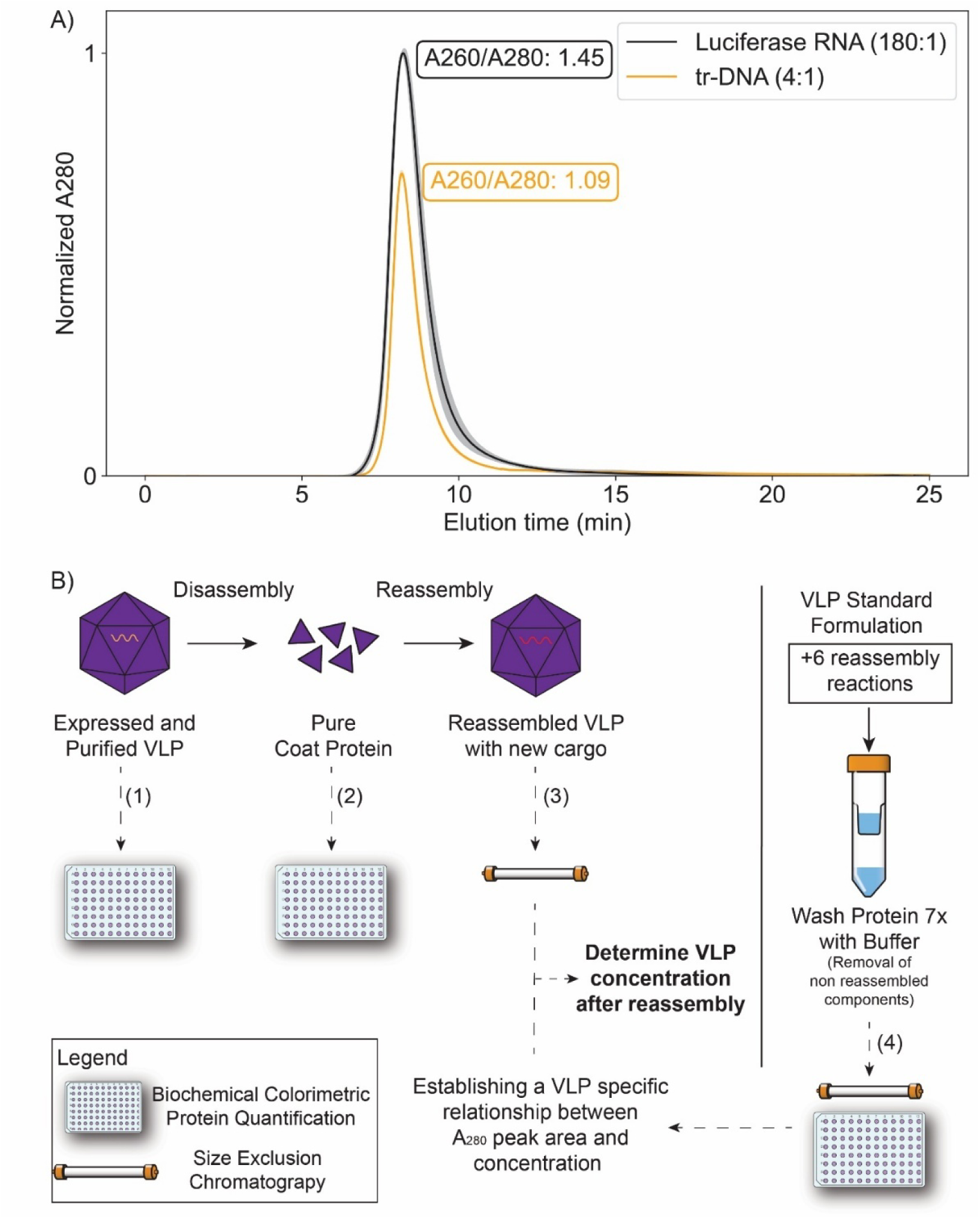
Standardized, cargo-based reassembly yield calculation method. A) SEC chromatograms normalized by A280 of two samples containing different nucleic acid-based cargos (varying in amount and nature) but with identical protein concentrations as measured by BCA. The resulting differences in absorbance highlight the interference caused by cargo. B) Schematic of the proposed method for quantifying VLP reassembly yield. The concentrations of the purified VLP (1) and the disassembled capsid (2) are determined using the BCA assay. The reassembly reaction product is analyzed by SEC (3). Additional reassembly reactions are performed to generate a standard curve relating SEC peak area to concentration (4). This standard is then used to convert the SEC peak area of the experimental reassembly product into concentration, enabling calculation of reassembly yield.

To continue to evaluate which protein concentration method is the most suitable, we assessed the sensitivity and linear dynamic range of BCA and SDS-PAGE methods. We also included UV-Vis based methods, either coupled to SEC or not, and using either the coupled A_280_ peak area or a NanoDrop^TM^ spectrophotometer, respectively. To this end, we expressed and purified MS2 VLPs to characterize with each method. We normalized the VLP-containing solution to A_280_ = 1, prepared dilutions spanning A_280_ values from 0.1 to 1.0, and measured each dilution using all four methods (**Supplemental Information 2**). Among them, NanoDrop^TM^ UV–vis spectroscopy, SEC, and BCA exhibited lower coefficients of variation between technical replicates compared to SDS-PAGE densitometry (average <u>c</u>oefficient of <u>v</u>ariance [CV] across nonzero datapoints: CV_UV–Vis_ = 0.02, CV_SEC_ = 0.008, CV_BCA_ = 0.03, CV_Densitometry_ = 0.1). Moreover, the relationship between expected A_280_ values and observed measurements revealed that densitometry of SDS–PAGE dilutions had a lower coefficient of determination and higher normalized root mean square error than the other methods (R²_Densitometry_ = 0.886, R²_BCA_ = 0.996, R²_UV–Vis_ = 0.999, R²_SEC_ = 0.999 | RMSE_Densitometry_ = 0.122, RMSE_BCA_ = 0.024, RMSE_UV–Vis_ = 0.012, RMSE_SEC_ = 0.011). These findings reinforce that densitometry-based methods suffer from intrinsically higher variability and a narrower linear dynamic range than absorbance-based techniques [22], rendering the common practice of single-point calibration (i.e., using a single band of known concentration to proportionally estimate the concentration of another band) unreliable. Nevertheless, accuracy limitations due to poor linearity can be mitigated by using a multi-point standard curve generated from purified coat protein and a high–dynamic-range gel imager. Finally, this dynamic range analysis indicates that the BCA assay does not span as broad a dynamic range as SEC coupled with A_280_ peak area. However, it remains the only tested method capable of reporting absolute protein concentration without interference from nucleic acids or other small molecules, making it a valuable complement to relative quantification approaches.

Given our findings, we propose a standardized and reproducible method for quantifying reassembly efficiency (**Figure 3B**) combining both separation of assembled and disassembled VLPs and yield quantification while addressing variability in how reassembly yield is reported in the literature. This approach allows for the calculation of either the reassembly yield alone (*Yr*) or the overall disassembly–reassembly yield (*Yg*), depending on which parameters are measured.

The core of this method involves generating a custom VLP standard specific to the VLP–cargo system under study. To do this, approximately six parallel reassembly reactions should be prepared using known amounts of coat protein and cargo to generate a study-specific VLP-cargo standard. After reassembly, these reactions should be pooled and subjected to centrifugation using a 100 kDa spin filter to remove unassembled components. The resulting purified sample represents fully reassembled VLPs that serve as the reference standard. This is then analyzed by both absorbance at 280 nm (in this case, measured by SEC) and BCA assay to determine protein concentration. From these two measurements, a conversion factor (*cf*) is calculated by correlating standard protein concentration (*C_S_*) determined by BCA with the A_280_ peak area (*A_S_*) measured by SEC (Equation 1):

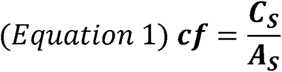

Because SEC has a broad linear dynamic range, a single calibration point—run in triplicate—is sufficient. However, for added robustness, performing a short dilution series and fitting a linear regression is recommended. This calibration allows the conversion of SEC A_280_ peak areas from any experimental reassembly sample (*A*) into capsid concentration (*C*) (Equation 2):

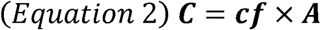

Since the standard and the reassembled samples share the same coat protein and cargo composition, the calibration factor derived from this standard is directly applicable. Once the capsid concentration of the reassembled sample (*C*) is determined, the reassembly yield (*Y_r_)* can be calculated using the concentration of disassembled coat protein used in the reassembly reaction (*C_CP_*), the volume of disassembled coat protein used in the reassembly (*V_CP_*), the volume of the reassembly reaction (*V_reassembly_*), and the volume injected in the chromatography system (V_injection_) – Equation 3:

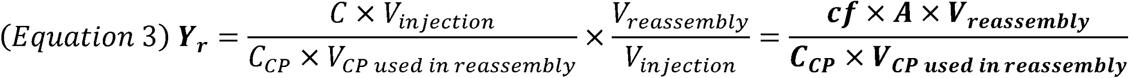

It is also important to consider the overall process efficiency, material losses during handling, and the practical feasibility of the method. To this end, we define the global disassembly–reassembly yield (*Yg*) metric, which quantifies the fraction of protein VLPs recovered relative to the initial amount subjected to disassembly. To calculate *Yg* (Equations 4 and 5), the following parameters are required: the concentration of VLPs prior to disassembly (*C_VLP_*), the volume of VLP solution subjected to disassembly (*V_VLP disassembled_*), and a correction factor to account for any volume changes introduced during post-disassembly purification (V*_CP stock after disassembly_* and V*_CP used in reassembly_*):

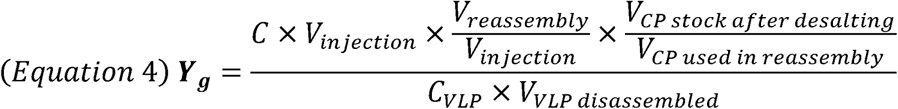

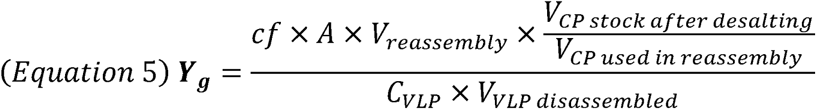

This standardized framework enables objective, reproducible, and cargo-specific quantification of VLP reassembly, improving the comparability of results across experiments and studies. With this method in hand, it becomes possible to systematically investigate how different variables influence the final reassembly yield.

### Impact of protein, crowding agent, sodium chloride concentration and buffer pH on VLP reassembly yield

Among the steps in the disassembly/reassembly process, the conditions used for reassembly vary the most. Coat protein concentration ranges from 0.05 μM of coat protein monomer [23] to 146 µM (2 mg/mL) [17]. The buffer pH ranges from 5.8 [23] to 8.5 [9], but is not always reported. The sodium content in the buffer ranges from 0 [9,13,21] to 100 mM [11,12], and some research groups add trimethylamine N-oxide (TMAO) at varying concentrations as a crowding agent [8,12]. The reassembly incubation temperature varies from 4 °C [8] to 48 °C [23]. Reported incubation time for the reassembly reaction varies from 10 min [10] to 51 h [16]. A summary of reported reassembly conditions and the corresponding publications can be found in **Supplemental Information 1.**

To understand the impact of these variables and their interactions on reassembly yield, we estimated the effect (quantitative influence that an independent variable has on a response) by performing a full factorial 2^4^ <u>d</u>esign <u>o</u>f <u>e</u>xperiments (DOE) with a central point, in which we combinatorially tested four factors (variables) with each one assuming two levels (values). The factors and levels we adopted in the combinatorial design of experiments are coat protein monomer molarity [10 μM, 30 μM], sodium chloride molarity [0 mM, 200 mM], TMAO molarity [0 mM, 250 mM], and buffer pH [5,7]. In addition to estimating the effects, we statistically modeled yield as a function of the tested experimental conditions to predict which conditions maximize yield. The proposed model is linear and accounts for all main effects of the independent variables (*x_1_, x_2_, x_3_, x_4_*) as well as their two- and three-way interactions.

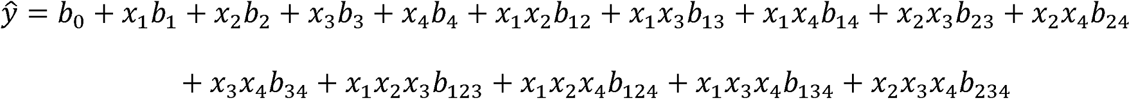

The modulus of the coefficients *b* indicates the intensity of the effect, and their signal indicates whether they have a positive or negative effect on yield. All the tested combinatorial conditions necessary for estimating the effects and generating the statistical model are represented in the experimental design matrix (**Supplemental Information 3**)

With the experimental results of the tested 17 conditions, we estimated the effects, their statistical significance (**Figure 4A**), the model parameters (**Table S3-5**), and standardized absolute effect (**Figure 4B**). Yield was quantified using the methodology developed in this study. Because the VLP standard for yield estimation was generated under the central-point condition (condition 17), yields for conditions with higher packing efficiency are likely overestimated and those with lower packing efficiency underestimated. However, this systematic bias is consistent across conditions and does not affect the interpretation of the comparative results. The interaction parameters protein-TMAO (*x1x2*), protein-TMAO-pH (*x1x2x4*), protein-NaCl-pH (*x1x3x4*) and TMAO-NaCl-pH (*x2x3x4*) are not statistically significant with a significance level α = 0.05 and were thus removed from the proposed model. We tested the model significance by performing an ANOVA, testing the null hypothesis H_0_ that all regression coefficients (except *b_0_*) are 0. The ratio between the regression mean square and the residual mean square (QM_R_ = 0.254802, QM_r_ = 0.000563, QM_R_/QM_r_ = 452.93) was higher than a critical F-value (ν_1_ = 10, ν_2_ = 40, α = 0.05, Fν_1,_ν_2_ = 2.077), and the ratio of the mean square for lack of fit and the mean square for pure error (QM_pe_ = 0.000514, QM_lof_ = 0.000838, QM_pe_/QM_lof_ = 1.632) was below a critical F-value (ν_1_ = 6, ν_2_ = 34, α = 0.05, Fν_1,_ν_2_ = 2.380). The results of the F-tests in conjunction with a model with coefficient of determination R^2^ = 0.9912 suggest that the model is statistically significant and adequately fits the data without significant lack of fit (**Figure 4C**).

**Figure 4:**
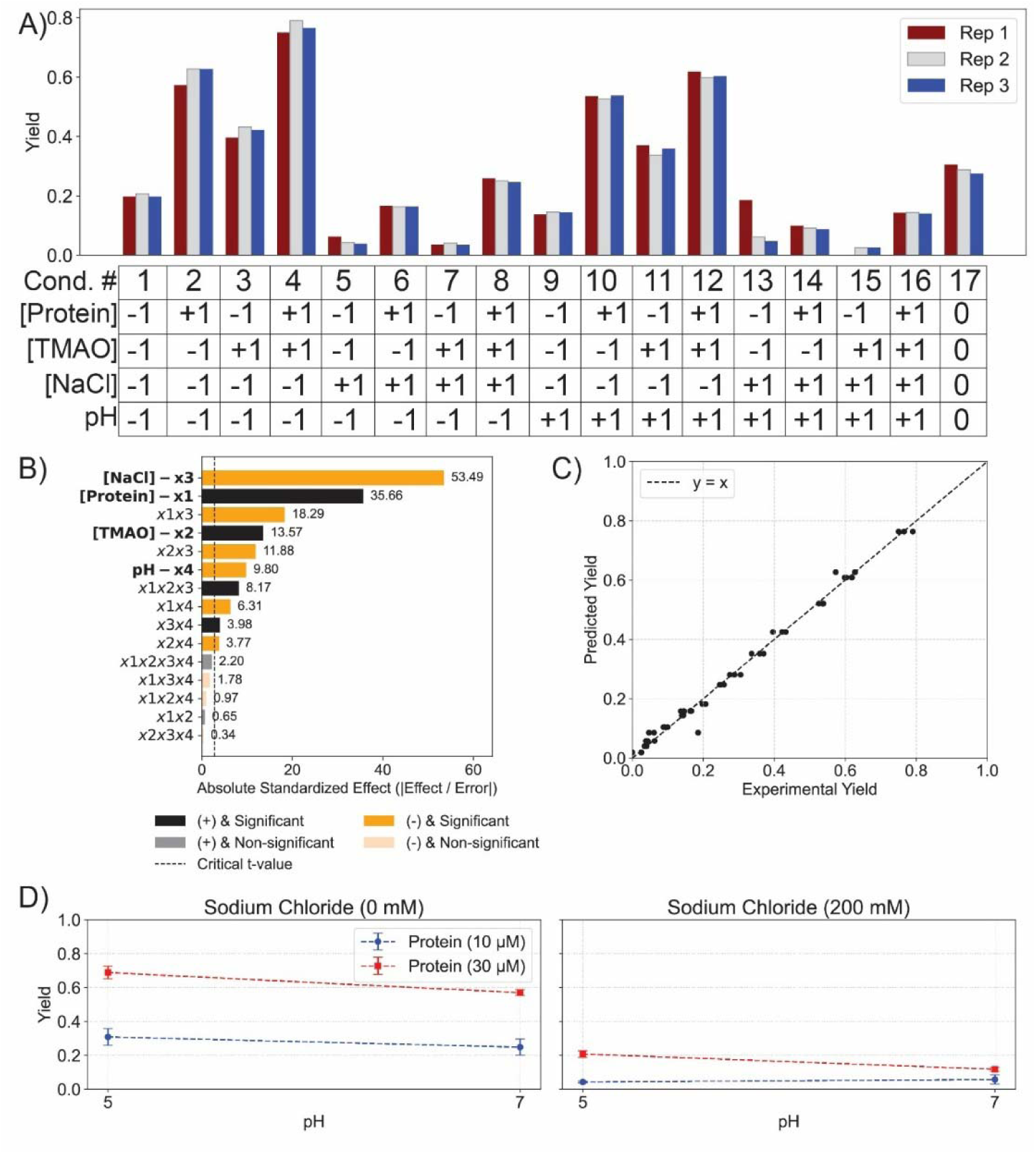
Determinants of Reassembly Efficiency. A) Reassembly yield for each individual replicate of each tested condition. B) Pareto chart of standardized effects showing the influence of individual factors and their interactions on yield. The vertical dashed line represents the critical t-value for statistical significance. C) Experimental versus predicted yield from the fitted model. Each point represents one replicate. The dashed line corresponds to the ideal fit (y = x), indicating strong agreement between experimental data and model predictions. D) Marginal means plot indicating the impact of protein molarity, sodium chloride molarity, and pH and their interactions on reassembly yield

The effect of protein concentration on the VLP reassembly yield is interconnected with other variables. While the main effect of protein concentration is one of the strongest in the model (0.232 ± 0.007, p-value = 3 ×10^-27^), it also had a strong negative interaction effect with sodium chloride (-0.1189 ± 0.007, p-value = 3 ×10^-15^) and a weaker negative interaction effect with pH (-0.0410 ± 0.007, p-value = 3 ×10^-6^). These interaction effects indicate that the effect of protein concentration is pronounced in certain conditions (*e.g.* pH 5, sodium chloride = 0 mM), but minimal in others (*e.g.* pH 7, sodium chloride = 200 mM) (**Figure 4D**). Conversely, sodium chloride had a consistently negative effect on reassembly, suggesting that 200 mM sodium chloride is too high of a concentration for efficient reassembly. We hypothesize that this is because elevated ionic strength disrupts the electrostatic interactions between coat proteins that are necessary for proper nucleation and propagation of capsid structures [24]. Interestingly, the magnitude of the interaction between protein concentration and sodium chloride concentration exceeds the independent effects of pH and TMAO, highlighting their comparatively greater impact on yield.

The effects of pH and the crowding agent TMAO on reassembly yield are more subtle. Within the ranges tested, these parameters produced a milder influence on reassembly yield, suggesting the reassembly process tolerates moderate pH shifts and osmolyte levels without negatively impacting reassembly yield, within the tested range. This finding complements the understanding of TMAO’s impact in MS2 VLP reassembly reported by our own research group. Glasgow *et al.* [12] reported that increasing TMAO concentration from 0 to 250 mM in reassembly reactions with 30 µM coat protein monomer, 3:5 protein monomer to cargo (tr-DNA) molar ratio, and 100 mM of sodium chloride increased the reassembly from approximately 10% to 50%. For the same coat protein monomer concentration (30 µM), sodium chloride concentration (100 mM), and pH (7), but for a coat protein monomer to tr-DNA cargo ratio of 2:1, our model predicts a reassembly yield variation from 34% to 38%. This difference suggests that the interaction between TMAO concentration and cargo concentration or protein to cargo ratio has a significant impact on reassembly yield. It is important to note that the method utilized to estimate yield in both studies is different, highlighting the necessity of the standardized method for measuring reassembly yield developed earlier in this manuscript.

Another important outcome of the model is the absence of curvature in the response surface, which indicates that the optimal conditions for reassembly are not within the range of conditions tested here. If an optimal set of conditions had been captured within the experimental space, we would expect to see a nonlinear model with maxima. Instead, the linear behavior observed suggests that the best conditions lie outside of the ranges reported in the literature and explored to date. While it would be possible to work with higher protein and TMAO concentrations, lowering the pH below that used in this work could result in buffer conditions close to those in which the reassembled capsid is not stable. While lowering the pH may be prohibitive, the DOE results imply that the field has not yet converged on optimal reassembly parameters, and that established conditions represent only partial, rather than maximal, efficiency. Another important observation is that these experiments were performed using tr-DNA as a cargo and physicochemical interactions between the tested conditions and other cargos may vary drastically. Therefore, extrapolating the results here to other MS2 VLP-cargo systems should be done carefully and additional studies with an expanded set of variables is necessary to determine optimal conditions. Importantly, while each new cargo may require its own optimization, the present findings highlight that not all parameters will need to be reinvestigated from the underlying basics. Overall, although perfect reassembly conditions remain elusive, we identify key parameters that help guide future efforts, and the quantitative yield framework introduced here finally provides a foundation for systematic, cargo-specific optimization going forward.

Based on the experimental results presented here, we propose a standardized workflow for MS2 VLP disassembly and reassembly with nucleic acid cargo to improve reproducibility and comparability across studies. Purified MS2 VLPs should be disassembled at high protein concentration (10 mg/mL) using a 2:1 (v/v) ratio of glacial acetic acid for 90 min at 4 °C, followed by removal of precipitated native RNA by centrifugation (≥17,000 × g, 15 min, 4 °C) and verification of RNA clearance by A_260_/A_280_ ≤ 0.65. Excess acid should then be removed via dialysis against 1 mM acetic acid without added salt. We recommend reassembling the VLP in a buffer at pH 5 with no sodium chloride and 250 mM of TMAO. The initial concentration of Coat Protein in a reassembly reaction should be close to 50 µM and the reaction should be carried at 4 °C for 48 h. Finally, reassembly yield should be quantified using cargo-specific SEC calibration curves anchored to absolute protein concentration measured by BCA. Adoption of this workflow provides a practical, experimentally grounded baseline for MS2 VLP reconstruction and enables systematic extension to alternative cargos and assembly conditions.

## CONCLUSION

In this study, we dissected the *in vitro* disassembly–reassembly process of MS2 VLPs to identify parameters that influence efficiency and reproducibility. We show that using a 2:1 acetic acid-to-VLP ratio at high protein concentrations with a 90-min incubation provides robust disassembly and RNA clearance. We also highlight the importance of selecting appropriate methods for buffer exchange, as trade-offs exist between protein recovery, accessibility, and processing time. Our comparison of quantification methods for reassembly yield reveals that electrophoresis and spectroscopy alone are insufficient for accurate determination; instead, we propose a standardized, cargo-specific calibration framework using HPLC-SEC coupled with BCA. Through factorial DOE analysis, we demonstrate that protein concentration is the dominant factor driving reassembly efficiency, whereas sodium chloride has a strongly negative effect, and pH and TMAO contribute more modestly. Notably, the absence of curvature in our response surfaces suggests that the optimal conditions for reassembly have not yet been defined within the literature and lie beyond the ranges typically reported.

These findings underscore two key points: (i) methodological variation has hindered progress toward reproducible, high-yield encapsulation, and (ii) the field would benefit from adopting standardized conditions and yield reporting practices. More broadly, while individual reassembly parameters have been explored empirically in prior studies, the field has largely lacked a quantitative approach capable of evaluating multiple variables and their interactions simultaneously. Although design of experiments (DOE) methodology has existed for decades, their application remains uncommon in the VLP assembly literature, where most studies vary one factor at a time. By implementing a statistically rigorous full factorial DOE, we show that reassembly yield is dominated by protein concentration and ionic strength, with comparatively modest contributions from osmolyte concentration under the tested conditions. Importantly, this analysis reveals interaction effects that are not accessible through one-variable-at-a-time approaches. These findings emphasize that a DOE-based approach is necessary to understand the cumulative and interacting effects of multiple physicochemical variables on reassembly yield.

We also acknowledge the limitations of this work, which include the focus on a single type of cargo (tr-DNA) and omission of other variables such as temperature, incubation time, or different cargo-to-coat protein ratios. Numerous studies have demonstrated successful MS2 VLP reassembly with a range of cargos, including diverse nucleic acids and proteins (summarized in **Supplemental Information 1**). However, even among studies employing the same cargos and reassembly conditions, reported yields vary substantially [12, 15], and in some cases differ by nearly twofold. This variability highlights the challenges associated with comparing results across literature and emphasizes the need for standardized experimental and yield-quantification frameworks. By introducing a cargo-specific, SEC-based calibration strategy anchored to absolute protein concentration, this work provides a common quantitative baseline that facilitates reproducibility, cross-study comparability, and rational troubleshooting of reassembly workflows. Future studies expanding these parameters and testing diverse cargos will be essential for developing broadly applicable guidelines. To that end, our study provides a systematic framework to improve reproducibility, comparability, and ultimately the reliability of MS2 VLPs as a versatile nanotechnology platform. While developed here for MS2 VLPs, this framework is readily extendable to other VLP systems that rely on disassembly–reassembly strategies.

## MATERIALS AND METHODS

### MS2 VLP expression

The strain used in this experiment is the same as that described by Robinson *et al* [25]. It consists of *E. coli* DH10B cells harboring the MS2 coat protein gene cloned into a pBAD33 vector. Key features of this plasmid include a chloramphenicol resistance cassette for selection and an arabinose-inducible pBAD promoter for controlled expression of the MS2 gene. Cells from a 30% glycerol stock were used to inoculate a 125 mL baffled shake flask containing 12 mL of autoclaved Lysogeny Broth (LB-Miller formulation) supplemented with 34 µg/mL chloramphenicol. Cultures were grown overnight (14–16 h) at 37 °C with shaking at 225 rpm. 10 mL of this overnight culture were used to inoculate a 2.8 L baffled shake flask containing 1 L of autoclaved LB-M medium and 1 mL of a 34 mg/mL chloramphenicol stock solution. The culture was incubated at 37 °C with shaking at 225 rpm until it reached an optical density at 600 nm (OD) of ∼0.6. At this point, VLP expression was induced by adding L(+)-arabinose to a final concentration of 0.02% (w/v). The culture was incubated for an additional 16–20 h at 37 °C and 225 rpm. Cells were harvested by centrifugation at 4,800 × *g* for 10 min at 4 °C, and the supernatant was discarded. The resulting pellet was placed on ice to halt further protein expression and resuspended in 50 mL of SEC buffer (10 mM sodium phosphate, 200 mM sodium chloride, pH 7.2). If not purified immediately, cell suspension was stored at −80 °C for later use.

### MS2 VLP purification

To recover and purify MS2 capsid proteins, the resuspended cell pellets were lysed using a probe sonicator (Fisher Scientific, Model FB120) set to 50% amplitude for 10 min (2-second pulses followed by 4-second intervals, for a total sonication program duration of 30 min), with samples kept on ice throughout sonication to prevent overheating. Lysates were clarified by centrifugation at 17,000 × *g* for 10 min at 4 °C, and the pellet containing cell debris was discarded. The resulting supernatant was mixed 1:1 (v/v) with saturated ammonium sulfate aqueous solution and incubated on a vibrating platform at 4 °C for at least 2 h to precipitate protein. Following incubation, samples were centrifuged again at 17,000 × *g* for 10 min at 4 °C. The supernatant was discarded, and the resulting pellet was resuspended in 3 mL of 1× SEC buffer. To remove residual ammonium sulfate, the sample was dialyzed overnight at 4 °C using a 7,000 molecular weight cut off (MWCO) (Snakeskin) dialysis membrane against 4 L of 1× SEC buffer. The dialyzed sample was clarified by centrifugation (21,000 × *g*, 5 min, 4 °C), followed by filtration through a 0.45 μm syringe filter. For purification, the sample was subjected to Fast Protein Liquid Chromatography (FPLC) using an ÄKTA Pure system (Cytiva) equipped with a HiPrep 16/60 Sephacryl S-500 HR column, using 1x SEC buffer as the mobile phase run at a flow rate of 1 mL/min at room temperature. Up to 5 mL of sample was injected per run, and MS2 capsid elution was monitored at A_280_, consistently appearing between 60 and 80 mL of elution volume.

### High-Performance Liquid Chromatography (HPLC)

VLP reassembly reactions were assessed using an HPLC system (Agilent 1260 infinity II) and an SEC column (Yara 4000, Phenomenex). The mobile phase consisted of 1 × SEC buffer (10 mM phosphate, 200 mM NaCl, pH 7.2). Runs were carried out at room temperature by injecting 100 µL of sample at the time with a total method length of 25 min. VLPs eluted between 7.7 and 9.0 min.

### Transmission Electron Microscopy

400-mesh copper grids with Formvar/carbon film (EMS P/N FCF400-Cu-50) were hydrophilized via glow discharge. We applied 10 μL of 150 nM purified capsid solution for 5 seconds, then blotted with filter paper. A 10 μL aliquot of 1% (w/v) uranyl acetate was added and immediately wicked away; this step was repeated once. A final 10 μL of stain was applied, incubated for 4 min, and then blotted. Grids were imaged on a JEOL 1400 Flash TEM with a Gatan OneView camera.

### Sodium Dodecyl Sulfate–Polyacrylamide Gel Electrophoresis (SDS-PAGE)

Proteins were resolved on 12.5% polyacrylamide gels prepared with a standard Laemmli formulation. Samples were mixed with Laemmli buffer containing 10% β-mercaptoethanol, heated at 95 °C for 5 min, and loaded alongside a molecular weight ladder (EZ-run, Fisher Scientific). Electrophoresis was carried out at 150 V for ∼1 h in 1× SDS running buffer until the dye front reached the bottom of the gel. Protein gels were scanned using Azure biosystems 600 and densitometry analysis was performed utilizing Fiji software (Schindelin et al., 2012).

### Bicinchoninic Acid Assay

Protein concentrations were determined using the bicinchoninic acid (BCA) assay (Pierce BCA Protein Assay Kit, Thermo Scientific) according to the manufacturer’s instructions, using a 96 well plate and BSA standards.

### MS2 VLP disassembly and reassembly

Native MS2 VLPs (≤1 week old) were concentrated to A_280_ > 25.0 (approximately 10 mg/mL) using a 100 kDa molecular weight cutoff centrifugal filter (Amicon Ultra-15, Sigma-Aldrich) spun at 3000 × *g* in a swinging-bucket rotor. Disassembly was initiated by adding glacial acetic acid at a 2:1 (v/v) acid-to-capsid ratio, followed by incubation on ice for 1.5 h. Samples were centrifuged at 17,000 × *g* for 15 min at 4°C to pellet residual nucleic acids. The clarified supernatant, containing disassembled capsids, was dialyzed at 4 °C against 1.5 L of chilled 1 mM acetic acid using a 3.5 kDa molecular weight cutoff dialysis cassette. Dialysis was performed in two steps: 3 h in fresh buffer, followed by overnight dialysis. Successful RNA removal was confirmed by an A_260_/A_280_ absorbance ratio of ∼0.6. Reassembly conditions were varied as noted in the results section. In all cases, water, buffer, salt, crowding agent, cargo, and protein were added at listed concentrations in the order listed here, and incubated for 48 h at 4 °C. The samples were centrifuged at 17,000 × g for 15 min and supernatant collected for further analysis.

## Abbreviations

ANOVA: Analysis of Variance
BCA: Bicinchoninic Acid Assay
CP: Coat Protein
DOE: Design of Experiments
HPLC: High Performance Liquid Chromatography
SDS-PAGE: Sodium Dodecyl Sulfate-Polyacrylamide Gel Electrophoresis
SEC: Size Exclusion Chromatography
TEM: Transmission Electron Microscopy
TMAO: Trimethylamine N-Oxide
UV-Vis: Ultraviolet-Visible light spectrum
VLP: Virus-Like Particle

## Supplementary material description

Supplemental Information 1 summarizes literature on MS2 VLP disassembly/reassembly conditions. Supplemental Information 2 compares linearity across protein quantification methods. Supplemental Information 3 provides supporting calculations for statistical design of experiments using reassembly yield as the response. Supplemental Information 4 contains raw data and results of statistical design of experiments using A_260_/A_280_ as the response.

## Acknowledgements

This work was supported by the National Science Foundation (CBET-2043973, to D.T.-E.). D.C.A. received support from the Chemistry of Life Processes Predoctoral Training Program at Northwestern University (NIH T32GM149439). M.M.M. was supported in part by the Northwestern University Graduate School Cluster in Biotechnology, Systems, and Synthetic Biology, which is affiliated with the Biotechnology Training Program. S.L. was partially supported by the Molecular Biophysics Training Program at Northwestern University (NIH 5T32GM140995-04). A.C.C. was supported by Conselho Nacional de Desenvolvimento Científico e Tecnológico (CNPq grant number 303063/2022-0). N.W.K. was supported by the U.S. Army Research Office (W52P1J-21-9-3023 P00003/CS-21-0701-001).

## Author’s Contributions (CRediT)

**D.C.A.:** Conceptualization (lead), methodology (lead), validation (lead), formal analysis (equal), investigation (equal), resources (supporting), data curation (lead), writing – original draft (lead); writing – review and edit (supporting), visualization (lead), project administration (equal), funding acquisition (supporting). **E.S.V.:** validation (supporting), formal analysis (supporting), investigation (equal), visualization (support). **M.M.M.:** investigation (supporting), data curation (supporting) **S.L.:** investigation (supporting) **C.E.M.:** writing – review and edit (equal) **A.C.C.:** formal analysis (equal), data curation (supporting) **N.W.K.:** writing – review and edit (equal), visualization (supporting) **D.T.-E.:** writing – review and edit (equal), project administration (equal), funding acquisition (lead).

