## Supplemental Information for "Process for Standardizing and Assessing the Parameters Governing MS2 Virus-Like Particle Reassembly around Nucleic Acid Cargo"

### Supplemental Information 1 – Literature summary of MS2 VLP disassembly and reassembly experimental conditions

| References | Initial MS2 Concentration | Acetic Acid to Protein Solution ratio (v/v) | Incubation time | Nucleic Acid Removal Method and conditions |
|---|---|---|---|---|
| Glasgow et al. (2012) | 10 mg/mL | 2:1 | 30 min | Centrifuge (4 °C, 16000 g, 20 min) |
| Ashley et al. (2011) | 10 mg/mL | 3 acetic acid to 1 protein (v/v) | 2 h | Centrifuged ("briefly") |
| Stockley et al. (2007) | cite Sugiyama & Hartman | 2:1 | 1 h | cite Sugiyama & Hartman |
| Sugiyama & Hartman (1967) | 1-2% | 2:1 | 1 h | Centrifuge ("in the cold", 8000 rpm, 20 min) |
| Garmann et al. (2019) | cite Sugiyama & Hartman | cite Sugiyama & Hartman | cite Sugiyama & Hartman | cite Sugiyama & Hartman |
| Zhang et al. (2015) | 4 mg/mL | 2:1 | 30 min | Centrifuge (4 °C, 6600 g, 20 min) |
| Wu et al. (1995) | 10 mg/mL | 2:1 | 30 min | Centrifuge (4 °C, 6600 g, 20 min) |
| Williams et al. (2024) | cite Sugiyama & Hartman | cite Sugiyama & Hartman | 30 min | Centrifuge (10000 g) |
| Aanei et al. (2018) | 10 mg/mL | 2:1 | 30 min | Centrifuge (4 °C, 14000 g, 10 min) |
| Yang et al. (2024) | n/a | 2:1 | 30 min | Centrifuge (4 °C, 12000 g, 30 min) |
| Zhou et al.(2025) | 10 mg/mL | 2:1 | 30 min | Centrifuge (4 °C, 10000 rpm, 20 min) |
| Rolfson et al. (2022) | n/a | cite Sugiyama & Nakada | cite Sugiyama & Nakada | cite Sugiyama & Nakada |

| References | Method for excess acidic acid removal |
|---|---|
| Glasgow et al. (2012) | Dessalting column (NAP-5) to 1 mM acetic acid |
| Ashley et al. (2011) | Dialysis (15 kDa MWCO tube) followed by dessalting column (Sephadex G-75). Dialysis buffer: 1.5 L (2 exchanges every 12 h) 10 mM acetic acid, 50 mM sodium chloride, pH 4. |
| Stockley et al. (2007) | cite Sugiyama & Hartman |
| Sugiyama & Hartman (1967) | Diluted protein/acetic acid solution to < 3 mg/mL. Dialyzed 3-4 times against 1 mM acetic acid at ice bath temperature. Centrifuge for 3 h at 40000 rpm |
| Garmann et al. (2019) | Centrifugal filtration (3 kDa MWCO), washing the protein with 20 mM acetic acid |
| Zhang et al. (2015) | Dialysis (7 kDa MWCO tube) followed by dessalting column (Superdex 75). Dialysis buffer: (3 exchanges every 4 h) 50 mM Tris, 100 mM sodium chloride. |
| Wu et al. (1995) | Dessalting column (NAP-25) to 1 mM acetic Acid |
| Williams et al. (2024) | Centrifugal filtration (3 kDa MWCO), washing the protein with 20 mM acetic acid (5 times) |
| Aanei et al. (2018) | Dessalting column to 1 mM acetic acid followed by centrifuge at 14000 g |
| Yang et al. (2024) | Dessalting column (5 mL) to 5 mM acetic Acid |
| Zhou et al.(2025) | Dessalting column (5 mL HiTrap) to 1 mM acetic Acid |
| Rolfson et al. (2022) | Exchange to 20 mM acetic Acid |

| References | Coat Protein Concentration | Cargo | Reassembly Buffer | Other agents |
|---|---|---|---|---|
| Glasgow et al. (2012) | 15 µM | yeast tRNA, tr-DNA, polyacrylic acid, mEGRP with and without 3xFLAG tag, PhoA-neg | Saline Tris Buffer (100 mM Sodium Chloride, 50 mM Tris) | TMAO - 250 mM |
| Ashley et al. (2011) | 1 mM of CP-dimers | siRNA, quantum dots, DOX | 50 mM Tris-HCl, pH 8.5 | |
| Stockley et al. (2007) | 8, 10, 20 µM CP-dimers | Synthetic oligonucleotides | 40 mM ammonium acetate, pH 6.8 | |
| Sugiyama & Hartman (1967) | | MS2 RNA | 0.1 M Tris-HCl buffer, pH 7.0 | |
| Garmann et al. (2019) | 1 - 4 µM CP dimers | MS2 RNA | Protein: 50 mM Tris-HCl, 100 mM NaCl, 1 mM EDTA / RNA: 10 mM Tris-HCl, 1 mM EDTA / | |
| Zhang et al. (2015) | 100 µM | DNA-*pac* site conjugates | 50 mM Tris with 100 mM NaCl | |
| Wu et al. (1995) | n/a | Deglycosylated ricin A chain conjugated to MS2 RNA stem-loop | 10 x TMK buffer (TMK, 100 mM Tris, 80 mM KCl, and 10 mM $MgCl_2$) | |
| Williams et al. (2024) | 2.5 - 30 µM CP-dimers | MS2 RNA | 42 mM Tris, 84 mM NaCl, 3 mM acetic acid, 1 mM EDTA, pH 7.5 | |
| Aanei et al. (2018) | 15 µM | GFP with encapsulation tags or gold nanoparticles | 50 mM bis-Tris, pH 6.0 | TMAO - 1.8 M |
| Yang et al. (2024) | 1 mg/mL dialyzed against 20 mM PBS and centrifuged at | none | 20 mM PBS | |
| Zhou et al.(2025) | 2 mg/mL | Protein nanocages | ST buffer (20 mM Tris-HCl, 100 mM NaCl), pH 7.2 | |
| Rolfson et al. (2022) | 0.1 - 7.5 µM CP-dimer | MS2 RNA fragment or smaller synthetic oligoribonucleotides containing tr stem loop | 20 mM Tris-acetate (pH 7.0-7.5) 40 mM ammonium acetate, 1 mM magneseium | |

| References | Incubation time | Incubation temperature | Methods used for yield calculation, if any |
|---|---|---|---|
| Glasgow et al. (2012) | 36 to 48 h | | Chromatography, using intact MS2 VLP as a standard |
| Ashley et al. (2011) | 1 h | Room temperature | SDS-PAGE |
| Stockley et al. (2007) | various times | | |
| Sugiyama & Hartman (1967) | various times, from 1 to 12 h | 37 C | Radioactively labeled coat protein and sucrose gradient centrifugation |
| Garmann et al. (2019) | Estimates capsid formation in about 300 s | | Native agarose gel electrophoresis |
| Zhang et al. (2015) | 43 to 51h | Room temperature (3h) then 4 °C (40-48h) | Size Exclusion Chromatography |
| Wu et al. (1995) | 36 h | Room temperature (3h) then 4 °C (33h) | |
| Williams et al. (2024) | 10 min | | Agarose gel and SDS-PAGE |
| Aanei et al. (2018) | 48 h | 4 °C | Area comparison in chromatography (capsid vs disassembled) |
| Yang et al. (2024) | | | |
| Zhou et al.(2025) | Overnight | | No |
| Rolfson et al. (2022) | 3 h | 48 °C | Agarose-acrylamide gels |

**Supplemental Information 2 – Linearity comparison among methods for protein quantification**

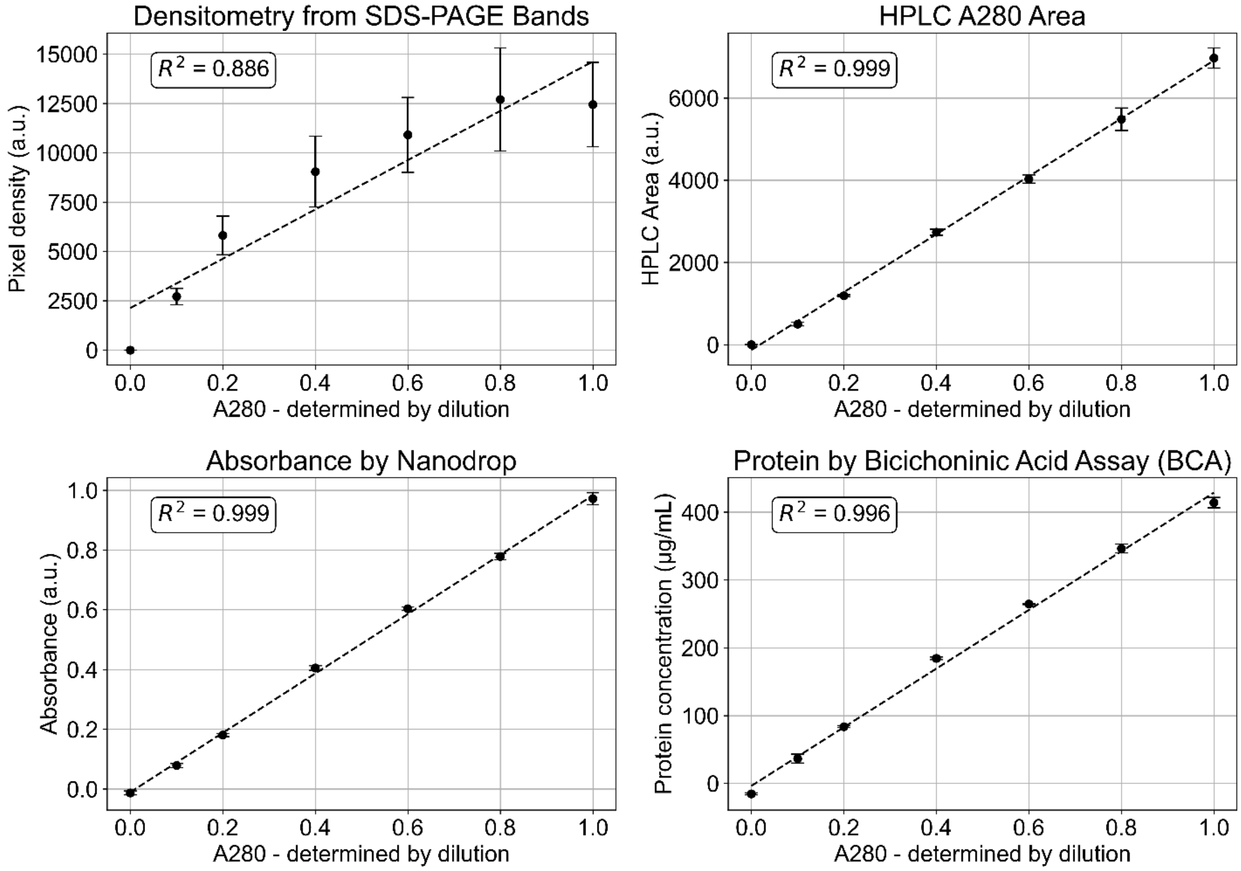


**Supplemental Figure 1:** Linear fits and respective coefficient of determination of data obtained from densitometry of SDS–PAGE bands, HPLC A280 peak area, UV–Vis spectroscopy, and BCA assay using pure MS2 VLP sample ($A_{280}$ = 1) and its dilutions.

**Supplemental Information 3 – Supporting calculations for statistical design of experiments using reassembly yield as the response.**

***Definition of experimental conditions and raw experimental data***

Using the design of experiment statistical methodology (Neto, Scarminio, & Bruns, 2010), we selected four independent variables for the reassembly reaction: coat protein molarity (µM, $X_1$), TMAO concentration (mM, $X_2$), sodium chloride concentration (mM, $X_3$), and pH ($X_4$) and created the experimental design matrix (**Table S3-1**):

**Table S3-1**: Experimental design matrix for assessing the effects of independent variables on VLP reassembly yield.

| ID | Protein Molarity (µM) - $X_1$ | TMAO (mM) - $X_2$ | Sodium Chloride (mM) - $X_3$ | pH - $X_4$ |
|---|---|---|---|---|
| **1** | 10 | 0 | 0 | 5 |
| **2** | 30 | 0 | 0 | 5 |
| **3** | 10 | 250 | 0 | 5 |
| **4** | 30 | 250 | 0 | 5 |
| **5** | 10 | 0 | 200 | 5 |
| **6** | 30 | 0 | 200 | 5 |
| **7** | 10 | 250 | 200 | 5 |
| **8** | 30 | 250 | 200 | 5 |
| **9** | 10 | 0 | 0 | 7 |
| **10** | 30 | 0 | 0 | 7 |
| **11** | 10 | 250 | 0 | 7 |
| **12** | 30 | 250 | 0 | 7 |
| **13** | 10 | 0 | 200 | 7 |
| **14** | 30 | 0 | 200 | 7 |
| **15** | 10 | 250 | 200 | 7 |
| **16** | 30 | 250 | 200 | 7 |
| **17** | 20 | 125 | 100 | 6 |

To enable statistical modeling, all variables were coded to a normalized scale of −1 to +1, resulting in a coded experimental design matrix (**Table S3-2**). The variable coding was done according to the transformation:

$$x_i = 2\frac{X_i - X_0}{X_i^+ - X_i^-}$$

where $x_i$ is the coded variable, $X_i$ is the uncoded experimental variable, $X_0$ is the center point, and $X_i^+$ and $X_i^-$ are the maximum and minimum values, respectively.

**Table S3-2:** Coded experimental design matrix for evaluating the effects of independent variables on VLP reassembly yield.

| ID | $x_1$ | $x_2$ | $x_3$ | $x_4$ |
|---|---|---|---|---|
| **1** | -1 | -1 | -1 | -1 |
| **2** | 1 | -1 | -1 | -1 |
| **3** | -1 | 1 | -1 | -1 |
| **4** | 1 | 1 | -1 | -1 |
| **5** | -1 | -1 | 1 | -1 |
| **6** | 1 | -1 | 1 | -1 |
| **7** | -1 | 1 | 1 | -1 |
| **8** | 1 | 1 | 1 | -1 |
| **9** | -1 | -1 | -1 | 1 |
| **10** | 1 | -1 | -1 | 1 |
| **11** | -1 | 1 | -1 | 1 |
| **12** | 1 | 1 | -1 | 1 |
| **13** | -1 | -1 | 1 | 1 |
| **14** | 1 | -1 | 1 | 1 |
| **15** | -1 | 1 | 1 | 1 |
| **16** | 1 | 1 | 1 | 1 |
| **17** | 0 | 0 | 0 | 0 |

Each condition of the experimental design matrix was performed in triplicate, and the area under the chromatographic curve (AUC) was measured at 280 and 260 nm (**Table S3-3**).

**Table S3-3:** Raw triplicate data obtained from the experimental design matrix.

| | HPLC A280 AUC (a.u.) | | | HPLC A260 AUC (a.u.) | | |
|---|---|---|---|---|---|---|
| **ID** | **1** | **2** | **3** | **1** | **2** | **3** |
| **1** | 206 | 210 | 207 | 263 | 274 | 262 |
| **2** | 1835 | 1994 | 1997 | 2314 | 2535 | 2533 |
| **3** | 426 | 466 | 455 | 525 | 573 | 559 |
| **4** | 2377 | 2517 | 2429 | 3030 | 3191 | 3092 |
| **5** | 77 | 55 | 50 | 83 | 58 | 51 |
| **6** | 648 | 633 | 652 | 673 | 659 | 663 |
| **7** | 44 | 49 | 43 | 47 | 54 | 46 |
| **8** | 1207 | 1206 | 1177 | 1044 | 1011 | 995 |
| **9** | 145 | 154 | 153 | 181 | 193 | 192 |
| **10** | 1724 | 1692 | 1724 | 2162 | 2123 | 2174 |
| **11** | 406 | 369 | 395 | 490 | 445 | 476 |
| **12** | 2031 | 1969 | 1992 | 2499 | 2416 | 2436 |
| **13** | 192 | 67 | 53 | 246 | 81 | 62 |
| **14** | 357 | 326 | 320 | 403 | 370 | 354 |
| **15** | 0 | 30 | 32 | 0 | 33 | 34 |
| **16** | 736 | 747 | 725 | 579 | 582 | 565 |
| **17** | 1051 | 991 | 947 | 828 | 781 | 746 |

***Calculation of reassembly yield and associated uncertainty***

To quantify reassembly yield, we first established a calibration factor ($c_f$) relating protein concentration measured by BCA ($C_S$) to the HPLC-SEC area under the curve ($A_S$). We created the protein standard discussed in the section "***Methods for determining Reassembly Yield and cargo impact on quantification***" and analyzed it using HPLC-SEC and BCA. We determined the conversion factor ($c_f$) between protein concentration obtained by BCA ($C_S = 806 \pm 51$ | average ± standard deviation) and area under the curve obtained in chromatography ($A_S = 5498 \pm 277$):

$$c_f = \frac{C_S}{A_S} = \frac{806}{5498} = 0.15 \frac{\mu g}{mL \times a.u.}$$

The reassembly yield (**Table S3-4**) for each replicate was then calculated as defined in the section "***Methods for determining Reassembly Yield and cargo impact on quantification***"

$$Y_r = \frac{c_f \times A \times V_T}{V_P \times C_{DP}}$$

where $V_T$ is the total reaction volume and $V_P$ the protein solution volume.

$$V_T = V_{Buffer} + V_{water} + V_{TMAO} + V_{Protein} + V_{NaCl} + V_{tr-DNA}$$

**Table S3-4:** Replicate reassembly yields, mean values, and associated errors for each condition, where i is the condition identification number (ranging from 1 to 17) and j is the replicate identification number (ranging from 1 to 3)

| i \ j | 1 | 2 | 3 | Average | Standard deviation |
|---|---|---|---|---|---|
| 1 | 0.1982 | 0.2068 | 0.1976 | 0.20087 | 0.0052 |
| 2 | 0.5731 | 0.6278 | 0.6273 | 0.6094 | 0.0314 |
| 3 | 0.3963 | 0.4324 | 0.4221 | 0.41694 | 0.0186 |
| 4 | 0.7503 | 0.7902 | 0.7657 | 0.76873 | 0.0201 |
| 5 | 0.0630 | 0.0435 | 0.0388 | 0.04842 | 0.0128 |
| 6 | 0.1666 | 0.1632 | 0.1642 | 0.16467 | 0.0017 |
| 7 | 0.0356 | 0.0405 | 0.0346 | 0.03688 | 0.0032 |
| 8 | 0.2585 | 0.2503 | 0.2464 | 0.25174 | 0.0062 |
| 9 | 0.1369 | 0.1459 | 0.1446 | 0.14247 | 0.0049 |
| 10 | 0.5356 | 0.5258 | 0.5385 | 0.53331 | 0.0066 |
| 11 | 0.3700 | 0.3362 | 0.3591 | 0.35506 | 0.0172 |
| 12 | 0.6189 | 0.5984 | 0.6033 | 0.60685 | 0.0107 |
| 13 | 0.1854 | 0.0615 | 0.0470 | 0.09797 | 0.0761 |
| 14 | 0.0997 | 0.0916 | 0.0877 | 0.09301 | 0.0061 |
| 15 | 0.0000 | 0.0250 | 0.0258 | 0.01694 | 0.0147 |
| 16 | 0.1434 | 0.1441 | 0.1400 | 0.14253 | 0.0022 |
| 17 | 0.3051 | 0.2878 | 0.2749 | 0.28929 | 0.0152 |

Using each condition variance ($s_i^2$), we estimated the overall experimental variance:

$$s^2 = \frac{\sum_{i=1}^{i=m} s_i^2 \nu_i}{\sum_{i=1}^{i=m} \nu_i} = 0.0005139$$

where m is the number of conditions (17) and each replicate has two degrees of freedom ($\nu_i = 2$).

***Estimating the effects of individual factors, 2-way, 3-way, and 4-way interactions and statistically modeling yield as a function of the experimental variables***

First, we define the contrast coefficient matrix (X). The matrix has 51 rows (17 conditions times 3 replicates). The first column consists of "ones", the second through fifth column corresponds to the individual factor levels and replicates as described in the experimental design matrix, and the subsequent columns represent interaction terms, calculated as the elementwise product of the corresponding independent factor columns (e.g., column $x_1x_2x_4$ is obtained by multiplying columns $x_1$, $x_2$, and $x_4$). Next, we define the response matrix (y), whose components are the individual values of each replicate for each tested condition.

| i | i | | Average | x1 | x2 | x3 | x4 | x1x2 | x1x3 | x1x4 | x2x3 | x2x4 | x3x4 | x1x2x3 | x1x2x4 | x1x3x4 | x2x3x4 | x1x2x3x4 | | |
|---|---|---|---|---|---|---|---|---|---|---|---|---|---|---|---|---|---|---|---|---|
| 1 | 1 | | 1 | -1 | -1 | -1 | -1 | 1 | 1 | 1 | 1 | 1 | 1 | -1 | -1 | -1 | -1 | 1 | | 0.19817 |
| 2 | 1 | | 1 | 1 | -1 | -1 | -1 | -1 | -1 | -1 | 1 | 1 | 1 | 1 | 1 | 1 | -1 | -1 | | 0.57313 |
| 3 | 1 | | 1 | -1 | 1 | -1 | -1 | -1 | 1 | 1 | -1 | -1 | 1 | 1 | 1 | -1 | 1 | -1 | | 0.39626 |
| 4 | 1 | | 1 | 1 | 1 | -1 | -1 | 1 | -1 | -1 | -1 | -1 | 1 | -1 | -1 | 1 | 1 | 1 | | 0.75031 |
| 5 | 1 | | 1 | -1 | -1 | 1 | -1 | 1 | -1 | 1 | -1 | 1 | -1 | 1 | -1 | 1 | 1 | -1 | | 0.06296 |
| 6 | 1 | | 1 | 1 | -1 | 1 | -1 | -1 | 1 | -1 | -1 | 1 | -1 | -1 | 1 | -1 | 1 | 1 | | 0.16657 |
| 7 | 1 | | 1 | -1 | 1 | 1 | -1 | -1 | -1 | 1 | 1 | -1 | -1 | -1 | 1 | 1 | -1 | 1 | | 0.03557 |
| 8 | 1 | | 1 | 1 | 1 | 1 | -1 | 1 | 1 | -1 | 1 | -1 | -1 | 1 | -1 | -1 | -1 | -1 | | 0.25852 |
| 9 | 1 | | 1 | -1 | -1 | -1 | 1 | 1 | 1 | -1 | 1 | -1 | -1 | -1 | 1 | 1 | 1 | -1 | | 0.13691 |
| 10 | 1 | | 1 | 1 | -1 | -1 | 1 | -1 | -1 | 1 | 1 | -1 | -1 | 1 | -1 | -1 | 1 | 1 | | 0.53556 |
| 11 | 1 | | 1 | -1 | 1 | -1 | 1 | -1 | 1 | -1 | -1 | 1 | -1 | 1 | -1 | 1 | -1 | 1 | | 0.36995 |
| 12 | 1 | | 1 | 1 | 1 | -1 | 1 | 1 | -1 | 1 | -1 | 1 | -1 | -1 | 1 | -1 | -1 | -1 | | 0.6189 |
| 13 | 1 | | 1 | -1 | -1 | 1 | 1 | 1 | -1 | -1 | -1 | -1 | 1 | 1 | 1 | -1 | -1 | 1 | | 0.18543 |
| 14 | 1 | | 1 | 1 | -1 | 1 | 1 | -1 | 1 | 1 | -1 | -1 | 1 | -1 | -1 | 1 | -1 | -1 | | 0.09969 |
| 15 | 1 | | 1 | -1 | 1 | 1 | 1 | -1 | -1 | -1 | 1 | 1 | 1 | -1 | -1 | -1 | 1 | -1 | | 0 |
| 16 | 1 | | 1 | 1 | 1 | 1 | 1 | 1 | 1 | 1 | 1 | 1 | 1 | 1 | 1 | 1 | 1 | 1 | | 0.14344 |
| 17 | 1 | | 1 | 0 | 0 | 0 | 0 | 0 | 0 | 0 | 0 | 0 | 0 | 0 | 0 | 0 | 0 | 0 | | 0.30515 |
| 1 | 2 | | 1 | -1 | -1 | -1 | -1 | 1 | 1 | 1 | 1 | 1 | 1 | -1 | -1 | -1 | -1 | 1 | | 0.20685 |
| 2 | 2 | | 1 | 1 | -1 | -1 | -1 | -1 | -1 | -1 | 1 | 1 | 1 | 1 | 1 | 1 | -1 | -1 | | 0.62777 |
| 3 | 2 | | 1 | -1 | 1 | -1 | -1 | -1 | 1 | 1 | -1 | -1 | 1 | 1 | 1 | -1 | 1 | -1 | | 0.4324 |
| 4 | 2 | | 1 | 1 | 1 | -1 | -1 | 1 | -1 | -1 | -1 | -1 | 1 | -1 | -1 | 1 | 1 | 1 | | 0.79021 |
| 5 | 2 | | 1 | -1 | -1 | 1 | -1 | 1 | -1 | 1 | -1 | 1 | -1 | 1 | -1 | 1 | 1 | -1 | | 0.04353 |
| 6 | 2 | | 1 | 1 | -1 | 1 | -1 | -1 | 1 | -1 | -1 | 1 | -1 | -1 | 1 | -1 | 1 | 1 | | 0.16318 |
| 7 | 2 | | 1 | -1 | 1 | 1 | -1 | -1 | -1 | 1 | 1 | -1 | -1 | -1 | 1 | 1 | -1 | 1 | | 0.04048 |
| 8 | 2 | | 1 | 1 | 1 | 1 | -1 | 1 | 1 | -1 | 1 | -1 | -1 | 1 | -1 | -1 | -1 | -1 | | 0.25031 |
| 9 | 2 | X = | 1 | -1 | -1 | -1 | 1 | 1 | 1 | -1 | 1 | -1 | -1 | -1 | 1 | 1 | 1 | -1 | y = | 0.14592 |
| 10 | 2 | | 1 | 1 | -1 | -1 | 1 | -1 | -1 | 1 | 1 | -1 | -1 | 1 | -1 | -1 | 1 | 1 | | 0.52584 |
| 11 | 2 | | 1 | -1 | 1 | -1 | 1 | -1 | 1 | -1 | -1 | 1 | -1 | 1 | -1 | 1 | -1 | 1 | | 0.33617 |
| 12 | 2 | | 1 | 1 | 1 | -1 | 1 | 1 | -1 | 1 | -1 | 1 | -1 | -1 | 1 | -1 | -1 | -1 | | 0.59836 |
| 13 | 2 | | 1 | -1 | -1 | 1 | 1 | 1 | -1 | -1 | -1 | -1 | 1 | 1 | 1 | -1 | -1 | 1 | | 0.06146 |
| 14 | 2 | | 1 | 1 | -1 | 1 | 1 | -1 | 1 | 1 | -1 | -1 | 1 | -1 | -1 | 1 | -1 | -1 | | 0.09164 |
| 15 | 2 | | 1 | -1 | 1 | 1 | 1 | -1 | -1 | -1 | 1 | 1 | 1 | -1 | -1 | -1 | 1 | -1 | | 0.02498 |
| 16 | 2 | | 1 | 1 | 1 | 1 | 1 | 1 | 1 | 1 | 1 | 1 | 1 | 1 | 1 | 1 | 1 | 1 | | 0.14415 |
| 17 | 2 | | 1 | 0 | 0 | 0 | 0 | 0 | 0 | 0 | 0 | 0 | 0 | 0 | 0 | 0 | 0 | 0 | | 0.28778 |
| 1 | 3 | | 1 | -1 | -1 | -1 | -1 | 1 | 1 | 1 | 1 | 1 | 1 | -1 | -1 | -1 | -1 | 1 | | 0.1976 |
| 2 | 3 | | 1 | 1 | -1 | -1 | -1 | -1 | -1 | -1 | 1 | 1 | 1 | 1 | 1 | 1 | -1 | -1 | | 0.62728 |
| 3 | 3 | | 1 | -1 | 1 | -1 | -1 | -1 | 1 | 1 | -1 | -1 | 1 | 1 | 1 | -1 | 1 | -1 | | 0.42215 |
| 4 | 3 | | 1 | 1 | 1 | -1 | -1 | 1 | -1 | -1 | -1 | -1 | 1 | -1 | -1 | 1 | 1 | 1 | | 0.76566 |
| 5 | 3 | | 1 | -1 | -1 | 1 | -1 | 1 | -1 | 1 | -1 | 1 | -1 | 1 | -1 | 1 | 1 | -1 | | 0.03879 |
| 6 | 3 | | 1 | 1 | -1 | 1 | -1 | -1 | 1 | -1 | -1 | 1 | -1 | -1 | 1 | -1 | 1 | 1 | | 0.16424 |
| 7 | 3 | | 1 | -1 | 1 | 1 | -1 | -1 | -1 | 1 | 1 | -1 | -1 | -1 | 1 | 1 | -1 | 1 | | 0.03459 |
| 8 | 3 | | 1 | 1 | 1 | 1 | -1 | 1 | 1 | -1 | 1 | -1 | -1 | 1 | -1 | -1 | -1 | -1 | | 0.24639 |
| 9 | 3 | | 1 | -1 | -1 | -1 | 1 | 1 | 1 | -1 | 1 | -1 | -1 | -1 | 1 | 1 | 1 | -1 | | 0.14457 |
| 10 | 3 | | 1 | 1 | -1 | -1 | 1 | -1 | -1 | 1 | 1 | -1 | -1 | 1 | -1 | -1 | 1 | 1 | | 0.53852 |
| 11 | 3 | | 1 | -1 | 1 | -1 | 1 | -1 | 1 | -1 | -1 | 1 | -1 | 1 | -1 | 1 | -1 | 1 | | 0.35905 |
| 12 | 3 | | 1 | 1 | 1 | -1 | 1 | 1 | -1 | 1 | -1 | 1 | -1 | -1 | 1 | -1 | -1 | -1 | | 0.60327 |
| 13 | 3 | | 1 | -1 | -1 | 1 | 1 | 1 | -1 | -1 | -1 | -1 | 1 | 1 | 1 | -1 | -1 | 1 | | 0.04703 |
| 14 | 3 | | 1 | 1 | -1 | 1 | 1 | -1 | 1 | 1 | -1 | -1 | 1 | -1 | -1 | 1 | -1 | -1 | | 0.08772 |
| 15 | 3 | | 1 | -1 | 1 | 1 | 1 | -1 | -1 | -1 | 1 | 1 | 1 | -1 | -1 | -1 | 1 | -1 | | 0.02583 |
| 16 | 3 | | 1 | 1 | 1 | 1 | 1 | 1 | 1 | 1 | 1 | 1 | 1 | 1 | 1 | 1 | 1 | 1 | | 0.14 |
| 17 | 3 | | 1 | 0 | 0 | 0 | 0 | 0 | 0 | 0 | 0 | 0 | 0 | 0 | 0 | 0 | 0 | 0 | | 0.27494 |

We assumed the response variable (ŷ) as a linear function of main effects, 2-way, and 3-way interactions:

$$\hat{y} = b_0 + x_1b_1 + x_2b_2 + x_3b_3 + x_4b_4 + x_1x_2b_{12} + x_1x_3b_{13} + x_1x_4b_{14} + x_2x_3b_{23} + x_2x_4b_{24} + x_3x_4b_{34} + x_1x_2x_3b_{123} + x_1x_2x_4b_{124} + x_1x_3x_4b_{134} + x_2x_3x_4b_{234}$$

Where $b_0$ is the intercept (overall mean response), $b_1$, $b_2$, $b_3$, $b_4$ are the coefficients of the main effects, $b_{ij}$ are the coefficients of two-factor interactions, $b_{ijk}$ are the coefficients of three-factor interactions, and $b_{1234}$ is the coefficient of the four-factor interaction. Each coefficient b quantifies the estimated contribution of the corresponding factor or interaction to the response variable. We estimated the model coefficients matrix ($b$) (**Table S3-5**):

$$b = (X^tX)^{-1}X^ty$$

and their respective variance matrix $s(b)^2$:

$$s(b)^2 = (X^tX)^{-1}s^2$$

where the error is determined as the non-zero value in the element-wise square root of the s(b).

We next calculated the respective effects which, by definition, are twice each model parameter (except for the average) and the effect error which is also twice the parameter error. Then, we tested the null hypothesis $H_0 : \beta = 0$ for each effect. The test statistics were computed as the ratio of the estimated effect to its standard error, and the corresponding p-values were obtained from a two-tailed Student's t distribution with 34 degrees of freedom ($\sum_{i=1}^{i=m} \nu_i$ with m = 17 and $\nu_i$ = 2). Effects with p-values greater than the significance level $\alpha = 0.05$ were considered statistically non-significant.

**Table S3-5:** Statistical significance of main and interaction effects of independent variables on MS2 reassembly, and the associated model coefficients (b). Values in red indicate p > 0.05 (non-significant).

| Parameter | Effect | s(effect) | t-value (effect/error) | p-value | b | s(b) |
|---|---|---|---|---|---|---|
| Average | 0.2809 | 0.0032 | 88.488 | $8.76{\times}10^{-42}$ | 0.2809 | 0.0032 |
| Protein monomer molarity (x1) | 0.2318 | 0.0065 | 35.427 | $3.01{\times}10^{-27}$ | 0.1159 | 0.0033 |
| TMAO molarity (x2) | 0.0882 | 0.0065 | 13.477 | $2.91{\times}10^{-14}$ | 0.0441 | 0.0033 |
| Sodium Chloride molarity (x3) | -0.3477 | 0.0065 | -53.130 | $1.16{\times}10^{-29}$ | -0.1738 | 0.0033 |
| pH (x4) | -0.0637 | 0.0065 | -9.733 | $3.71{\times}10^{-10}$ | -0.0318 | 0.0033 |
| x1x2 | 0.0042 | 0.0065 | 0.638 | $5.29{\times}10^{-1}$ | 0.0021 | 0.0033 |
| x1x3 | -0.1189 | 0.0065 | -18.170 | $9.80{\times}10^{-15}$ | -0.0595 | 0.0033 |
| x1x4 | -0.0410 | 0.0065 | -6.268 | $4.03{\times}10^{-6}$ | -0.0205 | 0.0033 |
| x2x3 | -0.0772 | 0.0065 | -11.795 | $6.65{\times}10^{-10}$ | -0.0386 | 0.0033 |
| x2x4 | -0.0245 | 0.0065 | -3.750 | $1.74{\times}10^{-3}$ | -0.0123 | 0.0033 |
| x3x4 | 0.0259 | 0.0065 | 3.954 | $1.44{\times}10^{-3}$ | 0.0129 | 0.0033 |
| x1x2x3 | 0.0531 | 0.0065 | 8.117 | $3.23{\times}10^{-6}$ | 0.0266 | 0.0033 |
| x1x2x4 | -0.0063 | 0.0065 | -0.962 | $3.58{\times}10^{-1}$ | -0.0031 | 0.0033 |
| x1x3x4 | -0.0116 | 0.0065 | -1.772 | $1.14{\times}10^{-1}$ | -0.0058 | 0.0033 |
| x2x3x4 | -0.0022 | 0.0065 | -0.340 | $7.45{\times}10^{-1}$ | -0.0011 | 0.0033 |
| x1x2x3x4 | 0.0143 | 0.0065 | 2.182 | $9.44{\times}10^{-2}$ | 0.0071 | 0.0033 |

Given the effects for the interactions $x_1x_2$, $x_1x_2x_4$, $x_1x_3x_4$, and $x_2x_3x_4$ are not statistically significant (**Table S3-5**), parameters $b_{12}$, $b_{124}$, $b_{134}$, and $b_{234}$ in the proposed model are also non-significant, thus, zero, resulting in the proposed model:

$$\hat{y} = 0.2809 + 0.1159x_1 + 0.0441x_2 - 0.1738x_3 - 0.0318x_4 - 0.0595x_1x_3 - 0.0205x_1x_4$$
$$- 0.0386x_2x_3 - 0.0123x_2x_4 + 0.0129x_3x_4 + 0.0266x_1x_2x_3$$

***Testing model significance using Analysis of Variance (ANOVA)***

Next, we performed an Analysis of Variance (ANOVA) (**Table S3-6**) and a variance F-test on the model.

**Table S3-6:** Equations used to calculate sums of squares, degrees of freedom, and mean squares for regression, residuals, lack of fit, and pure error. Also shown are equations for the coefficient of determination ($R^2$), maximum explainable variability, and the F-test used to evaluate model significance. Parameters: i = condition number (1–17); j = replicate number (1–3); $\bar{y}$ = overall mean of experimental measurements; $\hat{y}_i$ = model response for condition i; $\bar{y}i$ = mean of experimental measurements for condition i; $y_{ij}$ = experimental result for condition i, replicate j; p = number of model parameters; $n_i$ = number of replicates on condition i (3); n = total number of experimental (51); m = number of conditions (17).

| | **Quadratic Sum (QS)** | **Degrees of Freedom** | **Quadratic Mean (QM)** |
|---|---|---|---|
| Regression (R) | $\sum_{i}^{m}\sum_{j}^{n_i}(\hat{y}_i - \bar{y})^2$ | p – 1 | $\frac{QS_R}{p-1}$ |
| Residue (r) | $QS_{lof} + QS_{pe}$ | n – p | $\frac{QS_R}{n-p}$ |
| Lack of fit (lof) | $\sum_{i}^{m}\sum_{j}^{n_i}(\hat{y}_i - \bar{y}_i)^2$ | m - p | $\frac{QS_R}{m-p}$ |
| Pure Error (pe) | $\sum_{i}^{m}\sum_{j}^{n_i}\left(y_{ij} - \bar{y}_i\right)^2$ | n - m | $\frac{QS_R}{n-m}$ |
| Total (T) | $QS_r + QS_R$ | n – 1 | |
| Explained % of variability ($R^2$) | | $\frac{QS_R}{QS_T}$ | |
| Maximum Explainable % of Variability | | $1 - \frac{QS_{pe}}{QS_T}$ | |
| $F_{R,r}$ | $\frac{QM_R}{QM_r}$ | $F_{lof,pe}$ | $\frac{QM_{lof}}{QM_{pe}}$ |
| $F_{p-1,\, n-p}$ (α = 0.05) | | $F_{m-p,\, n-m}$ (α = 0.05) | |
| $F_{R,r}$ / $F_{p-1,\, n-p}$ | >10 | $F_{lof,pe}$ / $F_{m-p,\, n-m}$ | <1 |

The analysis (**Table S3-7**) decomposed the total variability (2.5705) into contributions from regression (2.5480) and residual error (0.0225). The residual error was decomposed into the model lack-of-fit (0.0050) and pure error (0.0175) contributions. The model explained 99.12% of the variability ($R^2$), from a maximum explainable variability of 99.32%.

To verify whether the model is significant, we computed the ratio between variance explained by the model ($QM_R$) and the variance explained due to random error ($QM_r$) ($QM_R$/ $QM_r$ = 452.93) and compared it to a critical value of $F_{10,40}$ = 2.077 (Fisher distribution with significance of 0.05). Given $F_{R,\,r} \gg F_{10,40}$, the model is statistically significant. To verify whether the model adequately fits the data or whether there is systematic deviation that the model is not capturing, we performed the F-test for lack of fit relative to pure error ($F = QM_{lof/pe} = 1.63$) was below the critical value ($F_{6,34} = 2.38$), indicating that the model adequately fits the data without significant lack of fit. These results demonstrate that the selected factors and interactions significantly account for the observed variability in the response.

**Table S3-7:** ANOVA results for the proposed model using the parameters on Table S3-6.

| Parameters: b0, b1. b2. b3. b4. b13. b14, b23, b24, b34, b123 | | | |
|---|---|---|---|
| | **Quadratic Sum** | **Degrees of Freedom** | **Quadratic Averages** |
| Regression (R) | 2.5480 | 10 | 0.254802 |
| Residue (r) | 0.0225 | 40 | 0.000563 |
| Lack of fit (lof) | 0.0050 | 6 | 0.000838 |
| Pure Error (pe) | 0.0175 | 34 | 0.000514 |
| Total | 2.5705 | 50 | |
| Explained % of variability ($R^2$) | | 0.9912 | |
| Maximum Explainable % of Variability | | 0.9932 | |
| $F_{Reg,res}$ | 452.93 | $F_{laf,pe}$ | 1.63 |
| $F_{10,40}$ ($\alpha = 0.05$) | 2.077 | $F_{6,34}$ ($\alpha = 0.05$) | 2.38 |
| $F_{R,r}/F_{10,40}$ | 218.04 | $F_{laf,pe}\backslash F_{7,15}$ | 0.69 |
